# A multi-tissue transcriptome analysis of human metabolites guides the interpretability of associations based on multi-SNP models for gene expression

**DOI:** 10.1101/773630

**Authors:** Anne Ndungu, Anthony Payne, Jason Torres, Martijn van de Bunt, Mark I. McCarthy

**Author notes:** These authors contributed equally to this work (Anne Ndungu, Anthony Payne, Jason Torres). These authors jointly supervised this work (Martijn van de Bunt, Mark I. McCarthy). OMNI Human Genetics, Genentech, 1 DNA Way, South San Francisco, CA 94080, USA.

## Abstract

There is particular interest in transcriptome-wide association studies (TWAS) - gene-level tests based on multi-SNP predictive models of gene expression - for identifying causal genes at loci associated with complex traits. However, interpretation of TWAS associations may be complicated by divergent effects of model SNPs on trait phenotype and gene expression. We developed an iterative modelling scheme for obtaining multi-SNP models of gene expression and applied this framework to generate expression models for 43 human tissues from the Genotype-Tissues Expression (GTEx) Project. We characterized the performance of single- and multi-SNP TWAS models for identifying causal genes in GWAS data for 46 circulating metabolites. We show that: (a) multi-SNP models captured more variation in expression than the top *cis*-eQTL (median 2 fold improvement); (b) predicted expression based on multi-SNP models was associated (FDR<0.01) with metabolite levels for 826 unique gene-metabolite pairs, but, after step-wise conditional analyses, 90% were dominated by a single eQTL SNP; (c) amongst the 35% of associations where a SNP in the expression model was a significant *cis*-eQTL and metabolomic-QTL (met-QTL), 92% demonstrated colocalization between these signals, but interpretation was often complicated by incomplete overlap of QTLs in multi-SNP models; (d) using a “truth” set of causal genes at 61 met-QTLs, the sensitivity was high (67%), but the positive predictive value was low, as only 8% of TWAS associations at met-QTLs involved true causal genes. These results guide the interpretation of TWAS and highlight the need for corroborative data to provide confident assignment of causality.

## Introduction

Genome wide association studies (GWAS) have been a powerful tool in revealing many loci that influence complex traits and diseases. However, most SNP associations map to non-coding regions of the genome, thereby complicating the task of identifying the (causal) genes through which the observed effects on disease predisposition are mediated^1^. To address this challenge, researchers have implemented a variety of approaches to link regulatory variants implicated in disease predisposition to their downstream effectors. One of the most widely adopted approaches leverages expression quantitative trait loci (eQTLs) to identify regional genes that are under the direct regulatory influence of the disease risk variant(s), and which thereby represent candidate mediators of disease predisposition. Empirical support for this approach is provided by the enrichment of *cis-*eQTL regulatory variants among significant GWAS variants and evidence that such variants explain a disproportionate share of trait heritability^2–6^.

A range of approaches have been deployed to detect coincident *cis-*eQTL and trait association signals. The simplest involves limiting the search space to trait variants that also demonstrate significant eQTL signals in a disease relevant tissue. In such analyses, it is now widely accepted that it is essential to test for statistical evidence of colocalization between eQTLs and trait-associated SNPs to avoid assigning relationships between eQTL and trait signals that map to distinct causal variants, and which cannot therefore be assumed to have any biological connection^7,8^.

Recently, this approach has been supplemented by a suite of methods (collectively, transcriptome-wide association studies or TWAS), built around a Mendelian randomization framework, which test for relationships between the genetic components of both complex traits and gene expression^5,9–13^. For example, the PrediXcan method generates predictive models of transcript expression from eQTL mapping data, and then uses these to “impute” estimates of gene expression into case-control or cohort-based GWAS datasets: those imputed estimates can then be subjected to trait association testing^12^. Although PrediXcan requires individual-level genotype data as input, there are conceptually similar approaches available that can accept GWAS summary statistics with linkage disequilibrium (LD) estimates from a reference population (e.g. S-PrediXcan, Fusion)^5,13,14^. Collectively, these methods have been applied to a broad range of complex traits and diseases and have spotlighted novel and biologically plausible candidate genes that had evaded detection in conventional GWAS approaches^5,11–13^.

The prediction models generated by these approaches range from those that feature only the single best (i.e. most strongly associated) eQTL for each gene, to those that support a polygenic model which comprises all SNPs within a locus (e.g. best linear unbiased predictor; BLUP). However, it has been shown that more sparse multivariate linear models (such as those generated by LASSO regression or a Bayesian Sparse Linear Mixed Model (BSLMM)) outperform both single variant and polygenic models in predicting gene expression^5,11–13,15^. Unlike single variant models, these sparse multi-SNP models can capture the effects of allelic heterogeneity (i.e. genes whose transcription is under the influence of multiple *cis-*regulatory signals). They also better reflect current understanding of the genetic architecture of gene expression than do polygenic models^5,12,15^.

The fact that multi-SNP models better predict gene expression than single-SNP models might suggest that trait associations based on these models would themselves involve multiple SNPs with shared effects on both expression and phenotype. However, the extent to which this is true is unknown. Moreover, if such models better reflect the number of independent genetic signals acting on a phenotype, are they supported by evidence of shared identity between the trait-associated and eQTL variants within the model? Furthermore, the extent to which novel genes implicated by colocalized associations represent genuine biological relationships, causal for disease, is unclear and inference is further complicated by the shared regulatory architecture of gene expression and by horizontal pleiotropy^16,17^.

To address these outstanding questions and guide the interpretability of predicted gene expression studies, we systematically evaluated sparse multi-SNP models underlying significant gene associations for evidence of independent effects on both phenotype and expression. We did this by generating multi-SNP gene expression models for 43 human tissues from the GTEx project and evaluating their utility through a large-scale analysis of GWAS data for 46 metabolites. We focused on metabolomic phenotypes as they provide a singular opportunity to assess the biological plausibility of multi-SNP gene associations. Insights from both genetic and experimental studies have led to well-curated sets of effector genes at loci with *cis-*associations to the levels of particular metabolites^18–21^. The subsets of genes so implicated encode enzymes, transporters, and regulators that can be directly tied to the specific metabolite, based on known functional relationships. These provide a “truth” gene set that can then be used to assess the performance (i.e. sensitivity and positive predictive value) of alternative analytical approaches for identifying effector transcripts, and which can inform the utility of applying TWAS approaches to the interpretation of GWAS data for other complex traits.

## Material and Methods

### GTEx expression data and *Cis*-eQTL analysis

Genotype data (variant call format), gene expression (quantified gene-level counts), and sample phenotype data from GTEx version 7 were obtained through dbGaP accession phs000424.v7.p2^22^. Genotypes were filtered to keep only bi-allelic variants with minor allele frequency of at least 0.05. Finally, only remaining SNPs that were tested in all metabolite GWAS were used for analyses to ensure consistent downstream modelling and testing across metabolites.

Only non-sex-specific tissue types with sample size of n ≥ 50 were analysed. For each tissue, genes reaching a threshold of > 6 raw reads and >1 count per million in at least 10 individuals were carried forward for analysis. Remaining genes were TMM normalised, then log transformed to counts per million using Voom^23^. Surrogate variables were calculated after explicitly defining sex in the models, and residual expression values after regressing out all surrogate variables and sex were used for analyses ^24^. Cis-eQTLs analysis was performed using QTLtools (Version 1.1) with a *cis-*distance limit of 1,000,000 base pair (1 Mb) from each gene^25^. The top eQTL SNP per gene was defined as the SNP with the lowest p-value for that gene.

### GWAS summary data

GWAS summary data for 46 metabolites were downloaded from the Metabolomics GWAS Server^20,26^. Metabolites for this analysis were selected based on having GWAS significant loci where at least one gene was identified as having a plausible or established biochemical link to the associated metabolite. Unknown metabolites and metabolite ratios were excluded from this analysis.

### LASSO regression, model filtering and final model selection

LASSO regression was used to select an optimal set of SNPs for explaining the expression of each gene. Regression was performed using GLMNET in R on each gene, with all SNPs less than 1MB from any part of each gene as potential covariates^27^. To select the optimal penalty factor for each gene, mean squared error (MSE) was calculated using 10-fold cross-validation across 100 automatically selected potential penalty factors. Given that data partitioning is random for cross-validation, this process was repeated 200 times per gene, and the penalty factor that had the mean lowest MSE across all iterations was selected as recommended in the reference manual for GLMNET.

For genes with multiple SNPs selected by LASSO regression, all selected SNPs were first linearly modelled against the gene’s expression. For any groups of SNPs in perfect LD, one was randomly selected and retained. Model *R*^2^ was calculated for the full linear model. Iteratively, starting with the SNP with the lowest p-value in the model, SNPs were added back one-at-a-time, each time calculating the subset model’s *R*^2^ (i.e. forward regression). Once 95% of the full model’s *R*^2^ value was attained; any SNPs not in the current subset model were eliminated. The final subset of SNPs was then modelled against expression and smoothed using ridge regression to minimize overfitting; with penalty factors selected using 25 iterations of 10-fold cross-validated ridge regression. For genes with only one SNP selected by LASSO, this SNP alone was modelled against gene expression using 25 iterations of 10-fold cross-validated ridge regression. The final coefficients from ridge regression models were carried forward for use in S-PrediXcan. Model fit p-values were determined by modelling pre-validated predicted expression of each gene against the observed expression. Model fit p-values were study wide FDR corrected (all genes and all tissues simultaneously), and those with adjusted p≥0.01 were excluded from further analysis due to poor model fit.

### Transcriptome wide association analysis with S-PrediXcan

For each modelled gene, S-PrediXcan (version 0.5.4) was used to calculate a z-score, which is a linear model of SNP effects for all SNPs in the gene’s final ridge regression model described above^14^. Each SNP’s effect is the product of its expression association coefficient, its GWAS z-score, and a SNP variance term (the SNP’s standard error divided by the standard error of the gene’s predicted expression). The SNP expression association coefficients used were those resulting from the final filtered gene expression ridge regression models. GWAS z-scores were calculated manually from effect size and standard error, since some SNPs had published p-values of 0 due to rounding.

### Conditional analysis

For significant genes identified by S-PrediXcan, we decomposed the z-scores into per-SNP scores. For each significant gene, for SNPs from the S-PrediXcan model that had the same individual direction of effect as the overall S-PrediXcan z-score, the SNP that had the highest absolute S-PrediXcan magnitude was considered the top contributing SNP for conditional analysis. Conditional analysis was performed on each significant S-PrediXcan gene using GCTA (version 1. 26.0)^28^. Each lead SNP effect was conditioned out of the GWAS summary data. S-PrediXcan was then performed as previously described, excluding the SNP/s being conditioned on, and using the GWAS z-scores resulting from the conditional GWAS analysis.

### Establishing biological plausibility of novel genes

Annotated protein information was downloaded from the Human Metabolome Database (version 3.6) on December 11, 2017^29^. HUGO gene names, metabolism pathways, and gene ontology classifications listed in this database were referenced to assess membership of significant S-PrediXcan associated genes. Metabolic pathways and GO classifications annotated to novel genes were compared with those for putative causal genes associated to the same metabolites to assess shared metabolic processes.

## Results

### Multi-SNP models explain more variance in gene expression than single eQTL models

To investigate gene associations based on multi-SNP models, we first evaluated the extent to which these models improve prediction of gene expression relative to single variant models. We obtained single variant models by performing standard univariate eQTL analysis to identify the top associated *cis*-SNP for each gene in each of 43 tissues from the GTEx study (version 7) with a sample size exceeding 50 (Methods) ^22^. The number of expressed genes (defined as genes with >6 raw reads and >1 count per million in at least 10 individuals), ranged from 15,483 in EBV-transformed lymphocytes to 19,846 in lung.

To obtain multi-SNP genetic prediction models of gene expression, we employed LASSO regression - a multivariate penalized regression procedure that simultaneously performs feature selection along with model fitting^27^ - to select an optimal and sparse set of *cis-*SNPs to jointly model expression of each gene in each tissue. We then compared the variation in gene expression explained by these multi-SNP models to that accounted for by the single eQTL models.

In Figure 1, we show representative results, in this case for skeletal muscle, the tissue with the largest sample size (n=385). LASSO regression selected multiple SNPs in the models for the majority of genes (n=11,210), and for these genes, there was a median of 2.4-fold increase (interquartile range or IQR, 1.7 to 3.9 fold) in expression variation explained by LASSO models versus the top eQTL alone (Figure 1A, B). There was a 2.0-fold median increase in expression variation explained across all gene models (i.e. including single-eQTL models) in skeletal muscle. LASSO selected the intercept-only model (i.e. model without any SNPs) for 2,667 genes out of 15,780 expressed genes, and the top-eQTL-only model (or a perfect proxy SNP) for 1,903 genes in skeletal muscle. The impact of multi-SNP selection seen for skeletal muscle was typical of that seen across all tissues and all genes (**Table S1**).

**Figure 1.**
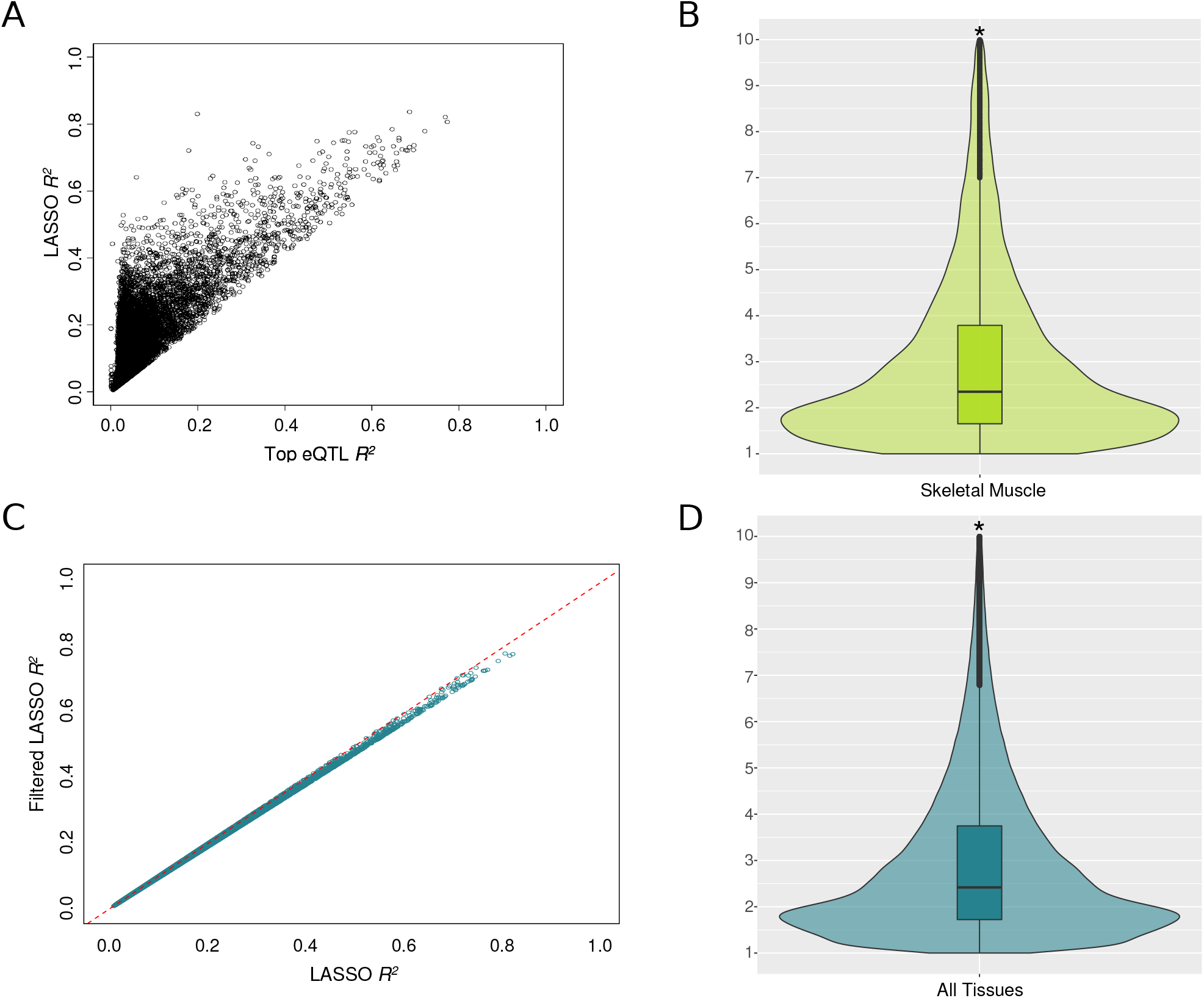
Model *R*^2^ comparison of LASSO regression models. **(A)** Scatterplot comparing variation in gene expression explained by the top eQTL alone and by the multi-SNP LASSO model in skeletal muscle. **(B)** Violin plot showing the fold increase in gene expression variation explained by LASSO models in skeletal muscle. The asterisks in the violin plots denote that the y-axis is abrogated at a fold change of 10. **(C)** Comparison of LASSO regression models before and after contribution based filtering of collinear SNPs. The mean model *R*^2^ was reduced by only 1.6%. **(D)** Violin plot showing the fold increase in gene expression variation explained by filtered LASSO models across all 43 tissues. The asterisks in the violin plots denote that the y-axis is abrogated at a fold change of 10.

Despite the sparse nature of LASSO selection, for those genes with at least two modelled variants, 7,406 genes (66%) in skeletal muscle retained at least one pair of SNPs with LD *r*^2^ >0.8. Moreover, LASSO expression models contained up to 159 SNPs and a median of 9 SNPs (IQR, 4 to 18 SNPs) for models with >1 SNP. Since correlated SNPs can result in invalid inference for summarised Mendelian randomisation (MR) analyses ^30^, we performed additional filtering of SNPs based on LD and proportion of variation explained (*R*^2^), iteratively adding SNPs into the model until 95% of the full model’s *R*^2^ was achieved. For groups of SNPs in perfect LD (*r*^2^ = 1), one SNP was randomly selected and retained (Methods). This reduced the median number of SNPs per gene in the model in skeletal muscle to six (IQR, 3 to 12 SNPs, **Table S1**). Moreover, 18% of gene models (2,015 out of 11,210 models that included multiple SNPs in the unfiltered analysis) contained only the top eQTL (or a perfect proxy). This further round of filtering had little impact on model performance; the mean reduction in model *R*^2^ was only 1.6% (calculated as percentages of the full LASSO models’ *R*^2^ values; Figure 1C,D). Similar to the unfiltered LASSO models in skeletal muscle, there was still a 2.0-fold median increase in expression variation explained across all gene models and across all tissues. Overall, these results demonstrate that multi-SNP models should generally be optimised and explained more variation in gene expression relative to the single top eQTL for the majority of genes across tissues.

### Transcriptome-wide association analysis of 46 metabolites across 43 tissues

Given these estimates of the extent to which multi-SNP models enhance the prediction of gene expression, we next sought to assess their utility in understanding genetic variation associated with complex diseases and traits. Metabolites offer a singular opportunity for such analyses as recent GWAS have identified strongly associated loci that regulate metabolite levels (met-QTLs) ^18–21^.

At some of these loci, extensive genetic and experimental evidence have identified nearby genes for which the biological evidence for a causal role in mediating the metabolomics association is overwhelming, providing a “truth” set for causal gene localization not available in most other trait GWAS settings.

We focused on 46 metabolites with publicly available GWAS data for which at least one gene mapped near a significant met-QTL signal with high confidence biochemical links to the associated metabolite (**Table S2**)^20^. We performed transcriptome-wide association analysis with S-PrediXcan^14^ to test for associations between predicted gene expression across 43 tissues and these 46 metabolite levels. Analysis was restricted to filtered LASSO prediction models with a strict significant expression model fit (model q<0.01; n=568,185 total gene models).

A total of 2,834 associations between predicted gene expression values and metabolite levels reached significance at study wide FDR <0.01, corresponding to 826 unique gene-metabolite pairs (i.e. some pairs were significantly associated in multiple tissues) (Figure 2A). The largest number of associations identified for any tissue was 100 (tibial nerve). There were only 66 associations arising from predictive models generated from liver expression data (8% of 826 unique associations), even though liver could be considered the most biologically relevant tissue for most of these metabolites. This is likely due to the relatively small sample size for liver in GTEx (153 samples compared to 361 in tibial nerve) (Figure 2B, Table S3).

**Figure 2.**
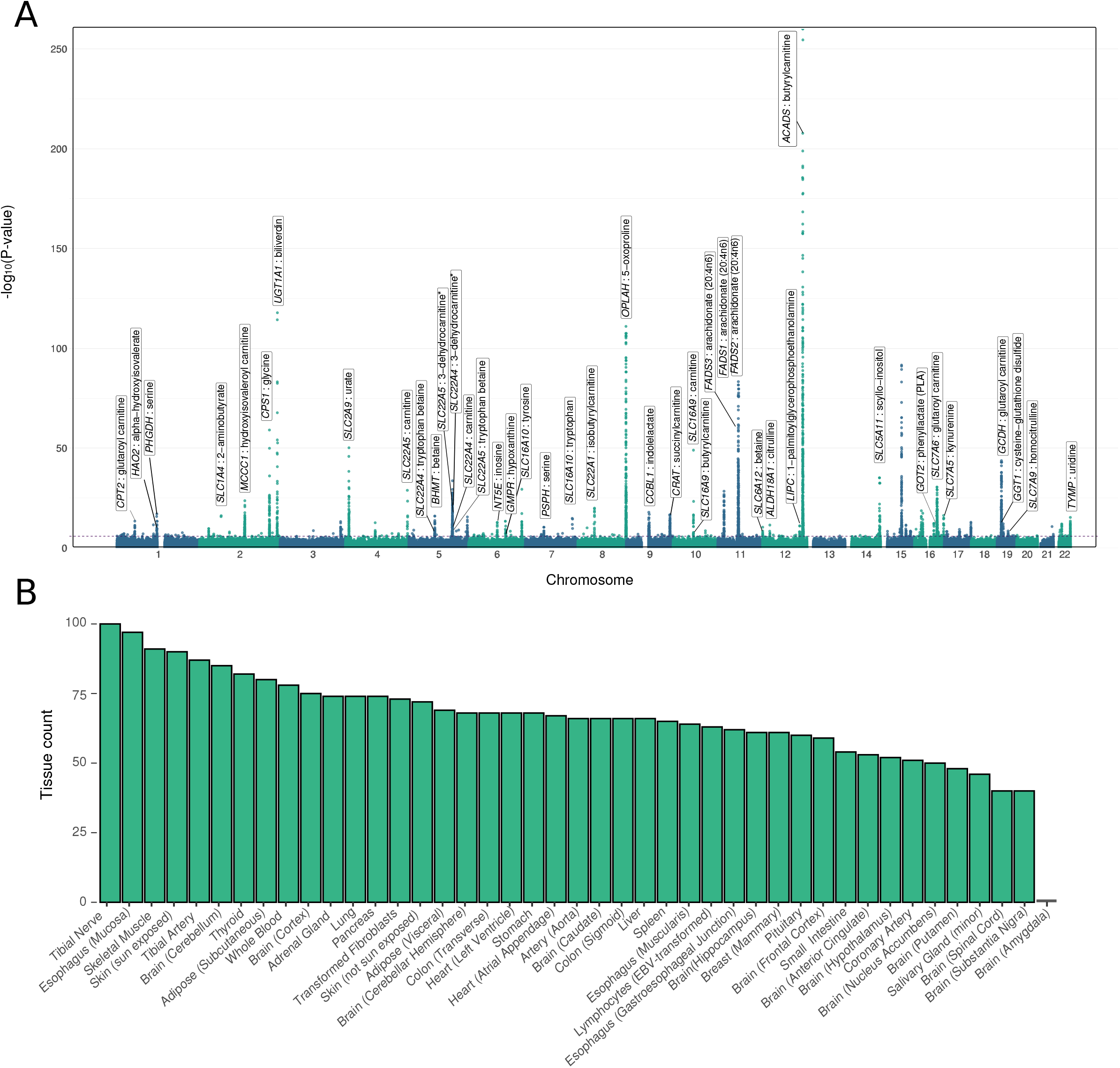
Transcriptome-wide association analysis of 46 metabolites across 43 tissues. **(A)** Manhattan plot showing all S-PrediXcan associations across 46 metabolites in all 43 tissues analysed with each point representing a gene-metabolite association. Labels indicate loci where TWAS associations involve high confidence causal genes. (**B)** Bar plot of the number of significant gene-metabolite associations observed per tissue.

For these 826 unique gene-metabolite pairs, we next sought to understand the extent to which multiple independent SNPs selected by the model were contributing to these metabolite associations. To do this, we performed conditional analyses for each of the 2,593 (from the total of 2,834) significant S-PrediXcan associations where the gene model had more than one SNP. We conditioned the metabolite GWAS on the SNP with the greatest effect on each gene’s S-PrediXcan score and re-ran the S-PrediXcan association test using the conditioned GWAS summary statistics. After correcting for the number of genes, tissues, and metabolites tested after conditional analysis (p-value _conditional_ <= 1.93×10^−5^), 2,320 of the 2,593 associations (89.5%) were no longer significant. This proportion was similar if we instead analysed only the most significant tissue for each gene; 684 out of 758 gene-metabolite pairs (90.2%) were no longer significant (p-value _conditional_ <= 6.61×10^−5^). Thus, for over 90% of significant S-PrediXcan associations, evidence for mediation of metabolite levels was dominated by a single SNP within the multi-SNP prediction models. Of the 273 still significant associations, over half (148) involved genes within 1 Mb of the highly complex *ACADS* gene region, which features multiple independent met-QTLs significantly associated with butyrylcarnitine levels (Figure 3, **Table S4**).

**Figure 3.**
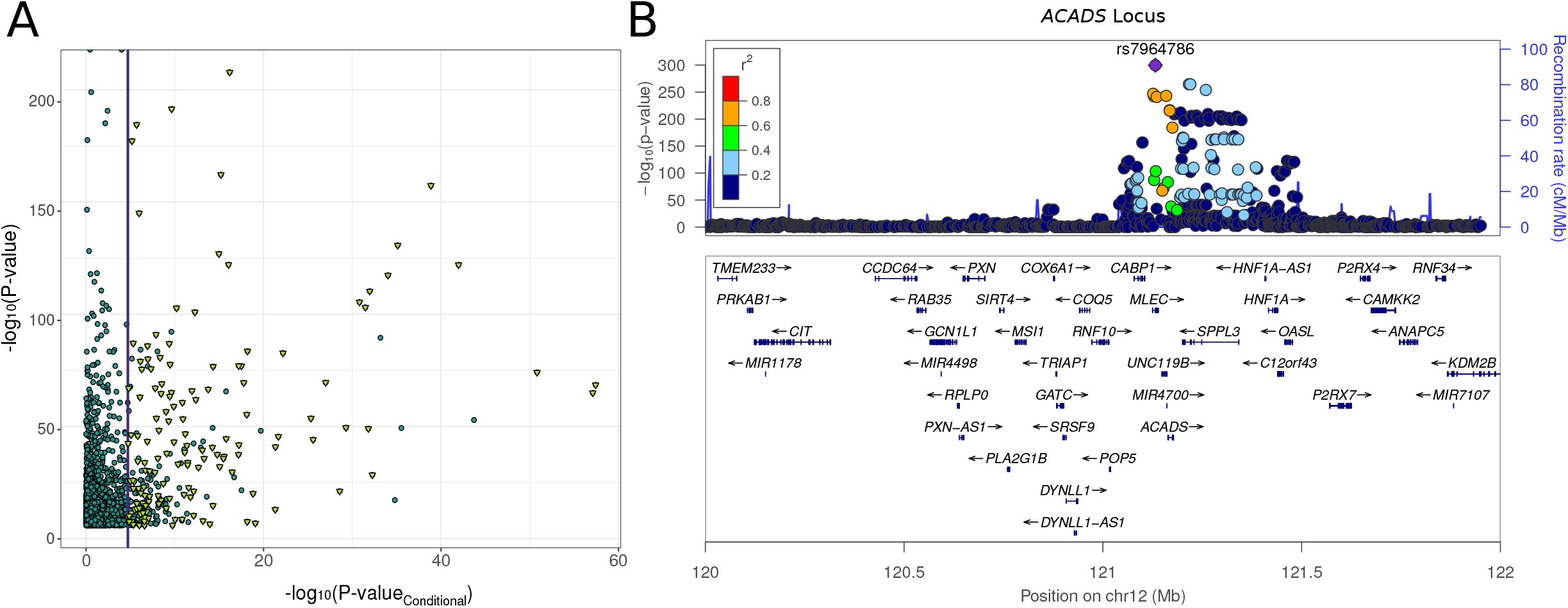
Step-wise conditional analysis of significant associations. **(A)** Plot showing results from the conditional analysis of S-PrediXcan associations involving multi-SNP prediction models. The vertical line denotes the significance threshold used for conditional analysis. Only 273 associations remained significant after conditioning on the lead met-QTL SNP, of which, 148 mapped to the *ACADS* locus and influence butyrylcarnitine levels (yellow triangles). **(B)** Locus Zoom plot showing met-QTLs associating with butyrylcarnitine levels at the *ACADS* locus and their LD relative to the top met-QTL.

### Colocalization analysis of model SNPs reveals the distinct relationships between *cis*-eQTL and met-QTL signals

It is possible that overlaps between GWAS met-QTLs and *cis-*eQTL variants in multi-SNP models could be due to chance, rather than representing true colocalization of causal signals. Consider, for example, a multi-SNP model with two SNPs where one SNP is a strong eQTL but weakly associated with metabolite levels, and the other SNP displays the converse arrangement: this configuration could still yield a significant association between gene expression and metabolite levels. We therefore questioned to what extent multi-SNP S-PrediXcan associations were driven by *cis-*eQTL and met-QTL signals that shared the same identity (i.e. the associations were attributable to SNPs that influence metabolite levels through their effects on gene expression).

We addressed this by performing colocalization analysis using eCAVIAR to obtain colocalization posterior probability (CLPP) values as evidence of shared causal signals, benefiting from the fact that eCAVIAR allows for multiple causal variants within a locus^8^. To increase our power to detect genuine colocalisation, we restricted this analysis to those SNPs in the prediction models that were significant *cis-*eQTLs (per tissue FDR<0.01) and met-QTLs (p-value<=5.0×10^−8^).

We found that, among the 2,834 significant S-PrediXcan associations, about 35% of associations (990 of 2,834 total; 214 unique gene-metabolite pairs) contained at least one SNP in the prediction model that influenced both metabolite levels at genome-wide significance and expression levels at FDR<0.05. Of these, 907 associations (92% of 990 associations tested; 202 unique gene-metabolite pairs) had at least one significant *cis-*eQTL with a CLPP > 0.01, evidence of a shared causal signal between met-QTL and *cis*-eQTL, in at least one tissue^8^ (**Table S5**). Therefore, for the SNPs that corresponded to gene models and that were amenable to colocalization analysis, there was strong evidence of shared eQTL and met-QTL signals.

We then analysed the context within which *cis-*eQTL SNPs in the multi-SNP models colocalized with met-QTLs. For the 907 associations with evidence of colocalization, we observed instances of a one-to-one overlap whereby the significant *cis-*eQTL in the multi-SNP model colocalized with the corresponding met-QTL. An example of this arrangement is displayed in Figure 4A. However, determining the evidence for or against colocalization of the met-QTL and *cis-*eQTLs was not always as simple, since many loci had a more complex topography. For example, expression of *SLC16A9* was significantly associated with carnitine levels in S-PrediXcan analyses in tibial nerve. Two significant *cis-*eQTLs with low LD (*r*^2^=0.002) were selected in the prediction model, but, as the locus plot shows, only one of these signals colocalized with the met-QTL (Figure 4B).

**Figure 4.**
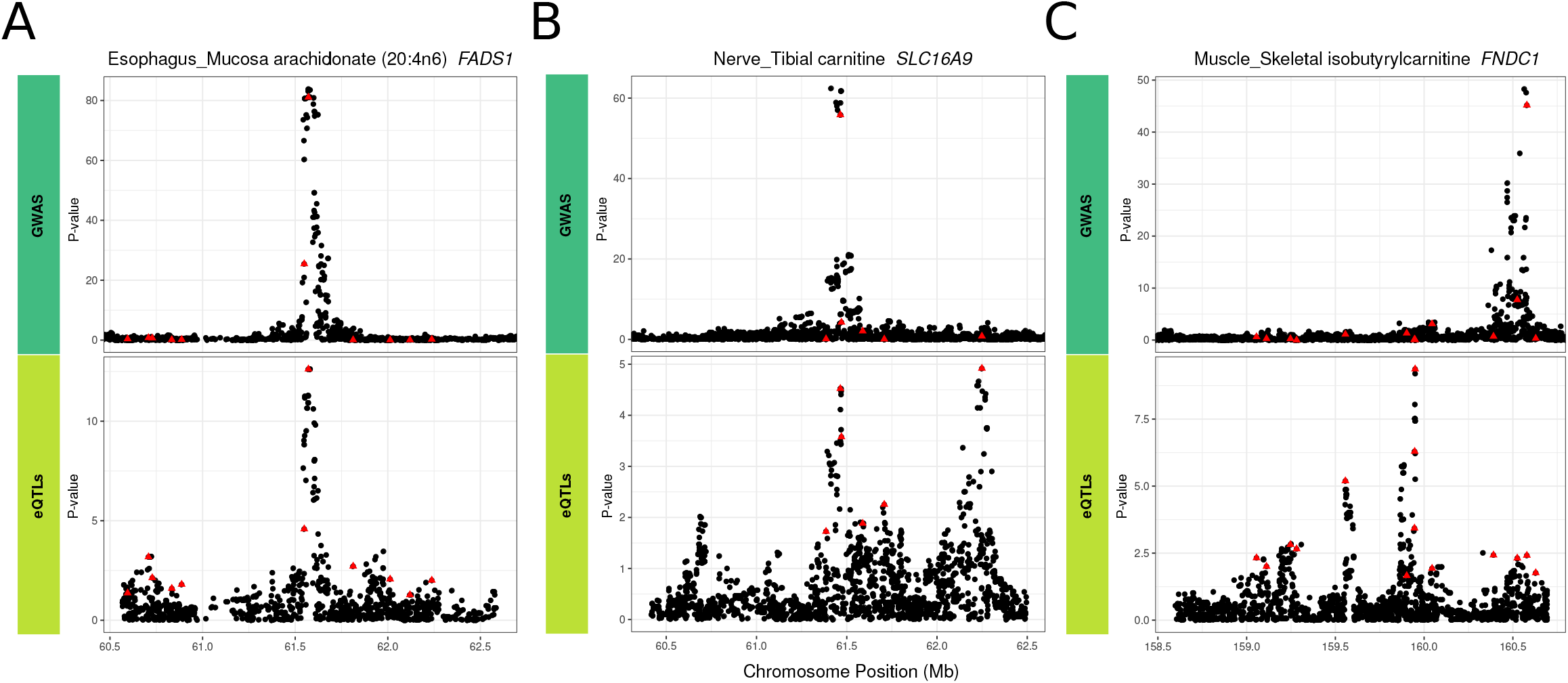
Colocalization analysis of eQTL and met-QTL signals in multi-SNP models for metabolite-associated genes. **(A)** Colocalization of the single met-QTL and single *cis-*eQTL signal at the *FADS1* gene in esophagus mucosa. **(B)** Partial colocalization at the *SLC16A9* gene in tibial nerve where only one of the two independent *cis*-eQTLs in the multi-SNP model is colocalized with the met-QTL at this gene (**C)** No colocalization of *cis-*eQTL and met-QTL for the *FNDC1* gene in skeletal muscle. The red triangles denote the SNPs present in the genes’ multi-SNP prediction models.

In contrast, we observed significant TWAS associations where model SNPs had divergent effects on expression and metabolite levels and were thereby excluded from colocalization analysis (i.e. associations not included in the 907 associations with evidence of colocalized QTL signals). For example, the expression of *FNDC1* in skeletal muscle was significantly associated with circulating isobutyrylcarnitine levels. However, the met-QTL and *cis-*eQTL were clearly not colocalized even though the genetically predicted expression of *FNDC1* was significantly associated with metabolite levels. This is because the set of SNPs in the *FNDC1* prediction model includes both the SNP driving the strong met-QTL (which explains a small portion of the variance in *FNDC1* expression) and a strong *cis-*eQTL that is only weakly associated with metabolite levels (Figure 4C).

### Determining the sensitivity and positive predictive value of multi-SNP prediction models

Across the 46 metabolite GWAS that we used as the substrate for our analyses, Shin et al. previously reported 61 SNP-metabolite associations at which the associated met-QTL SNP mapped near a gene that was highly likely to be causal for the association. This assessment was based on either experimental validation or a strong biological plausibility that the encoded protein was involved in the synthesis or degradation of the metabolite concerned^20^. These 61 SNP-gene-metabolite groupings provide a “truth” set of causal genes that can be used to explore the performance of expression QTL based mapping strategies, information relevant to more common applications (e.g. in a disease GWAS) where the causal gene is typically not known with equivalent certainty.

Of these 61 gene-metabolite pairs in the “truth” set, we were able to detect 41 through significant S-PrediXcan associations in at least one GTEx tissue (Table 1), indicating a sensitivity for *cis*-eQTL validation of the causal gene of 67%. Thirty-three of these gene-metabolite assignments were supported in more than one tissue with the *GCDH-*glutarylcarnitine association being the most widely represented (detected in 38 tissues, Table 1). Only eight of the 41 were detected in liver, though this may in part reflect the relatively small sample size of liver in GTEx. We assessed the extent to which overlaps between eQTLs and GWAS at these truth set genes represented true colocalization of signals. Of these 41 genes, 23 were amenable to colocalization analysis (i.e. at least one of the SNPs in the model was a significant *cis-*eQTL and a significant met-QTL) and all of these 23 genes showed evidence of colocalization, where at least one SNP in the multi-SNP model colocalised with the met-QTL in at least one tissue.

**Table 1.**
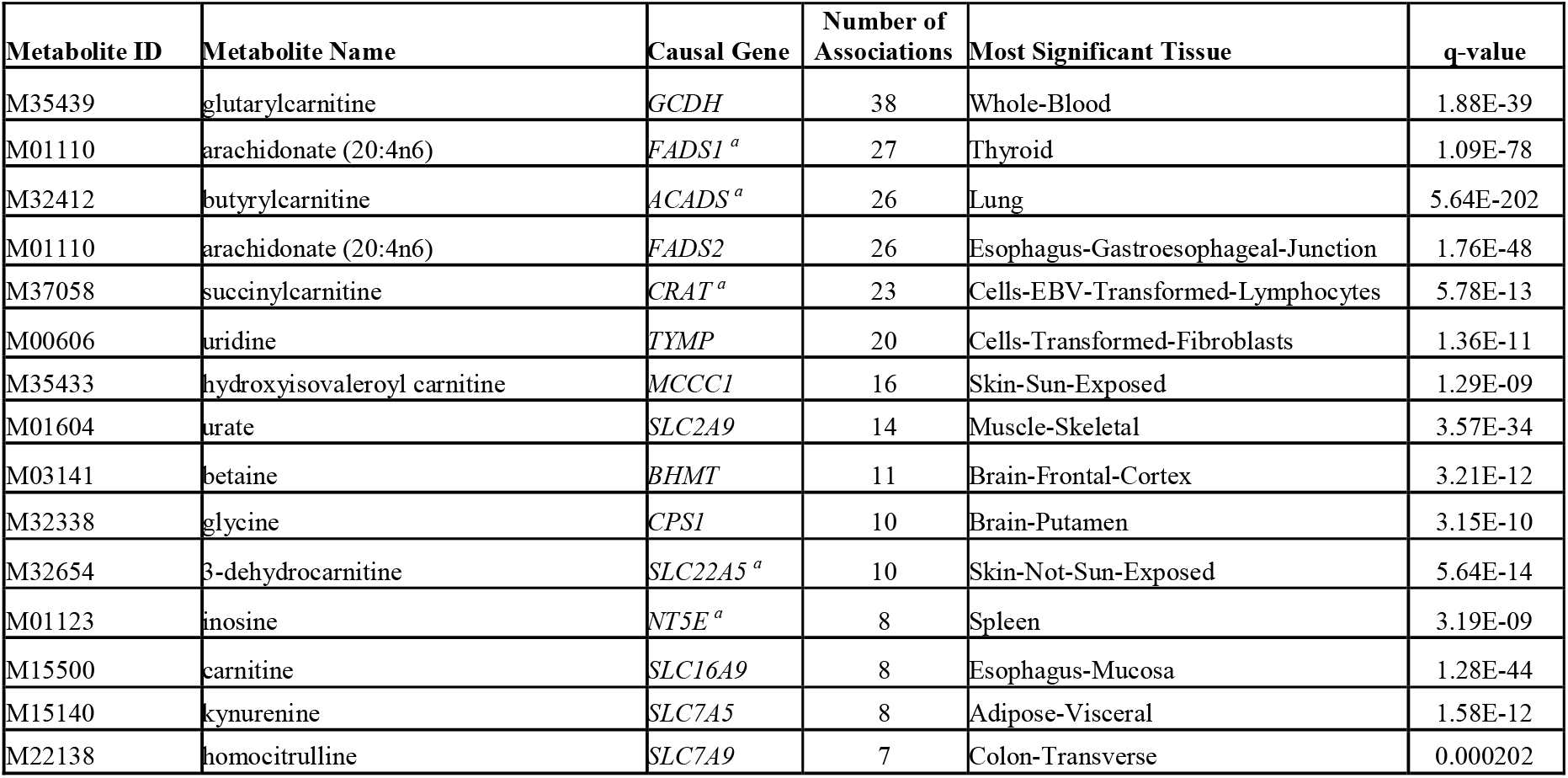
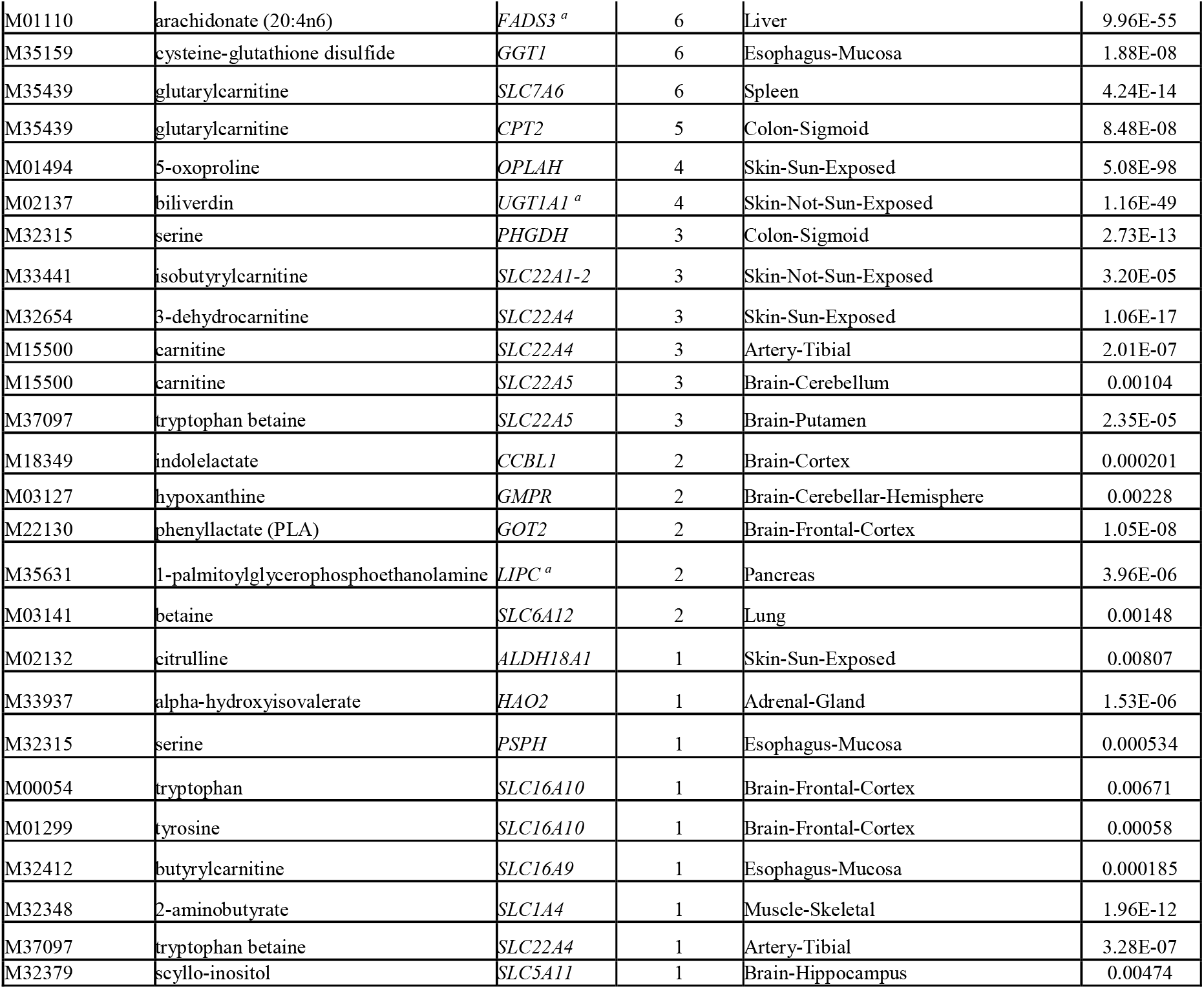
Causal genes from the truth set that significantly associated with metabolite levels in a TWAS. Of the 61 high confidence truth set genes, 41 had significant S-PrediXcan associations in at least one tissue. ^**a**^ Eight gene-metabolite pairs that had a significant association in liver.

As described earlier, our genome-wide trawl for associations between metabolite levels and predicted expression levels across GTEx tissues had implicated 826 unique gene-metabolite pairs. Of these, more than half (514; 62%) involved genes that mapped within 1Mb of the 61 truth set genes (including the 41 detected truth set genes). At only four of the truth set loci did these analyses identify the true causal gene only with no nearby bystander gene. This indicates that, at many of these loci, there are multiple “bystander” genes, other than the truth set genes, that are also being detected through predicted expression. From this analysis of TWAS associations at metabolite-associated loci, we estimate that the positive predictive value (PPV; i.e. number of true positives divided by the sum of true and false positives) for detecting true positive associations is only 8% (41/514 gene-metabolite pairs).

There were 20 of the 61 gene-metabolite pairs in the truth set that did not yield significant S-PrediXcan associations in any tissue. However, for 15 of these, significant S-PrediXcan associations (from the set of 514 gene-metabolite pairs described above) were seen for nearby bystander genes in at least one tissue, with eight of these showing significant bystander gene colocalization (**Table S4**). Taken together with the results for the 41 true positive signals, these analyses indicate substantial pleiotropy at the level of *cis-*eQTLs, with many met-QTL loci harbouring a substantial excess of “bystander” genes alongside the true causal gene (or at some loci, only “bystander” genes).

To illustrate these concepts, consider SNP rs8012, which is a significant met-QTL for glutarylcarnitine levels (p-value_GWAS_=1.24×10^−43^), and maps 8kb from the *GCDH* gene that encodes the enzyme glutaryl-CoA dehydrogenase. This enzyme catalyses the conversion of glutaryl-CoA to crotonyl-CoA, making *GCDH* a highly plausible effector gene mediating the effects of rs8012 on glutarylcarnitine levels^31^. In GTEx, whilst rs8012 is a *cis-*eQTL for *GCDH* in 31 tissues, the same SNP is also associated with the expression of other nearby genes including *HOOK2, SYCE2, FARSA, AD000092.3* and *CALR*. For all these genes, the *cis-*eQTL and the met-QTL signal clearly colocalized in at least one tissue (**Figure S1).** In the absence of the strong biological prior favouring *GCDH* at this locus, there would be at least five other genes that could be equally plausible candidates.

We next asked whether there were any features of the 473 bystander genes that might allow them to be distinguished from truth set genes. We found that bystander genes did not differ with respect to the strength of association with the metabolite, distance to transcription start site, the effect sizes of the individual eQTLs included in the multi-SNP models, or the CLPP values for model SNPs (Figure 5). However, we did find that causal genes tended to be significant in more tissues than bystander genes at the same locus (**Figure S2)**.

**Figure 5.**
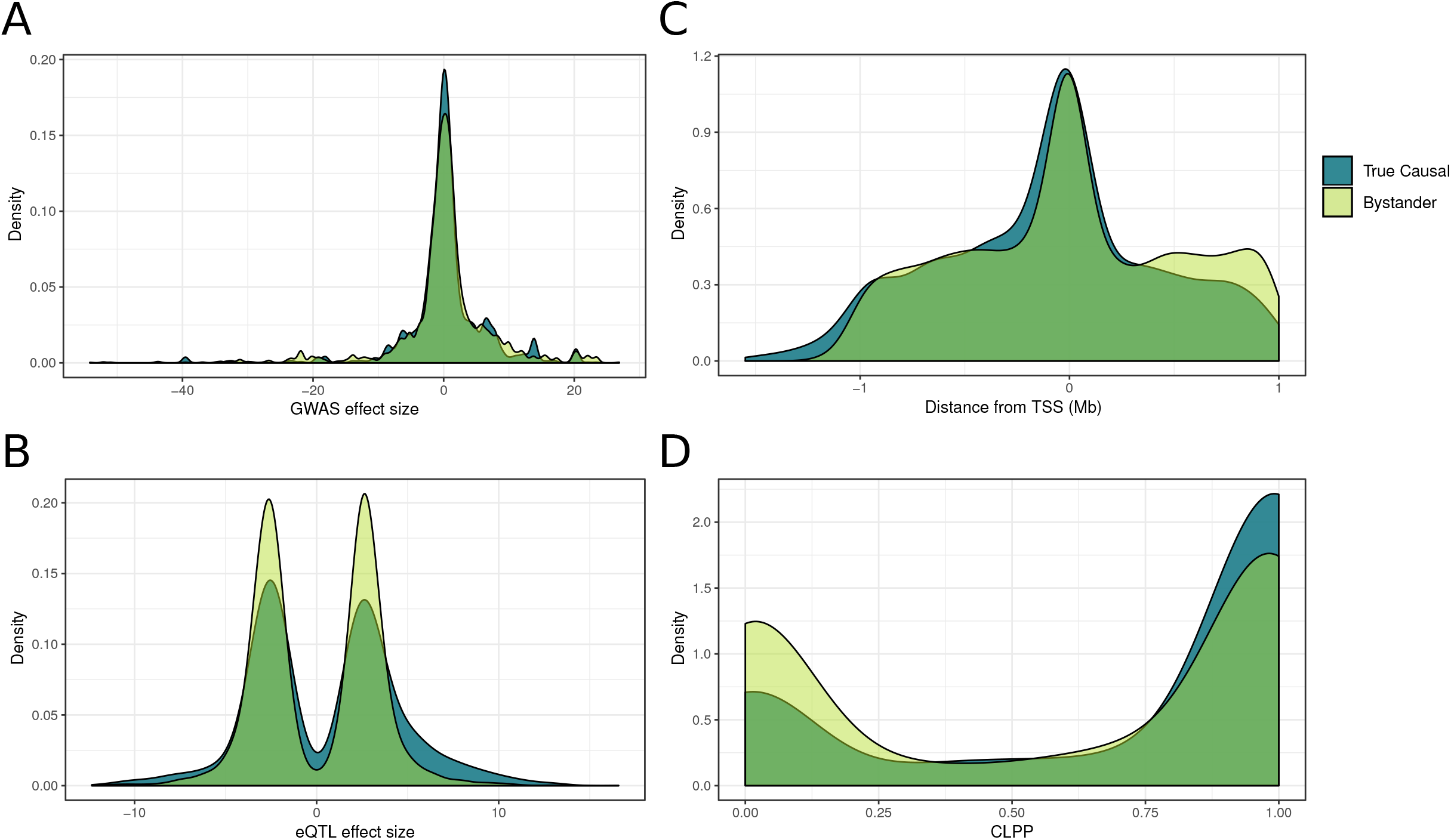
Comparison of features of multi-SNP models for bystander genes to those for true causal genes. **(A)** Comparison of the effect sizes of model SNPs for bystander genes and model SNPs for true causal genes on metabolite levels in GWAS. **(B)** Distribution of effects on gene expression for individual SNPs in models for bystander and known causal genes. (**C)** Comparison of the distance from TSS for model SNPs in bystander and causal genes. **(D)** The distribution of colocalization posterior probabilities (CLPP) scores for model SNPs in bystander and causal genes.

In addition to the 61 SNP-metabolite pairs in the truth set, Shin et al. reported 18 SNP-metabolite pairs that reached genome-wide significance in their analysis, but for which it was not possible to assign a causal gene with high confidence, as none of the genes could be implicated based on known biology^20^. In this setting, the authors assigned each associated SNP to the nearest gene at the locus (**Table S2)**. The results of our analyses for these 18 signals recapitulated those we saw for the 61 genes in the “truth set”. We could recover 10 of these “nearest gene” candidates (a sensitivity of 56%) using S-PrediXcan applied to GTEx (**Table S6**), of which seven colocalized in at least one tissue. However, a further 92 bystander associations at these loci were also significant.

We also used a complementary approach to quantify the performance of the predicted expression analysis for identifying novel, plausible gene candidates. We focused on the 312 gene-metabolite pairs that involved genes that did not map to known met-QTL regions and evaluated metabolite and gene annotations in the Human Metabolome Database (version 4.0)^29^. We found that 96 of these pairs - corresponding to 83 genes - involved genes annotated to metabolic pathways. These included two genes involved in uridine metabolism - *CDA* and *UPP1*. Notably, SNPs in at these two loci were sub-genome- wide significant in the GWAS but were implicated from our S-PrediXcan analysis and subsequent studies^17^ (**Table S7)**. We expanded the search further by querying a recently curated dataset^17^ and found an additional 18 genes annotated to at least one metabolic pathway. Thus, as many as 37% (114/312 gene-metabolite pairs) of novel TWAS gene associations can be considered biologically plausible albeit based on the rather permissive overlap between “metabolic pathway” and met-QTL.

We then performed a more stringent evaluation by determining the number of novel gene-metabolite associations (again excluding “bystander” genes) where the novel gene either shared at least one metabolic pathway with a reported truth set gene for the associated metabolite or has been curated as a high confidence causal gene with the associated metabolite in recent publications^17^. We found that 16 (5%) of the 312 novel gene-metabolite pairs met this criterion **(Table S8)**. Taking this as a lower limit and the previous less stringent estimate as an upper limit, we estimate that 5-37% of novel gene-metabolite relationships are biologically plausible. Notably, this range encompasses our PPV estimate of 8% obtained from evaluating the true positive rate at met-QTLs with known causal genes. Therefore, most novel gene associations based on multi-SNP models represented either false positives or “bystander” genes that are not biologically relevant *per se* but rather driven by variants with pleiotropic effects on gene expression. Overall, these findings emphasize that, whilst the multi-SNP *cis-*eQTL approach has respectable sensitivity in detecting the causal gene in these data, performance in terms of PPV is poor and additional lines of evidence will be needed at most loci to establish causality.

## Discussion

In this study, we have assessed the utility of multi-SNP prediction models for explaining variation in gene expression and their application in transcriptome-wide association analysis (TWAS). We quantified the extent to which these models outperform expression models based on a single eQTL, demonstrating, across all evaluated tissues, a median 2-fold improvement in variance explained. When applied in a TWAS of genome-wide data for 46 metabolites across 43 human tissues, these multi-SNP models identified 826 significant gene-metabolite associations. By leveraging knowledge of genes highly likely to be causally involved in the regulation of metabolite levels, we were able to quantify the accuracy with which multi-SNP TWAS detects such high-confidence effectors. The results from these analyses offer several key insights relevant to the interpretation of TWAS results.

We found that, notwithstanding the use of LASSO regression as a sparse form of variable selection, it is still prone to select sets of SNPs that are highly correlated, introducing multicollinearity into resulting regression models. This notion has been described before in real and simulated GWAS data^32^. We showed that a simple iterative approach to LASSO modelling that involved LD-based filtering resulted in increased model sparsity and decreased multicollinearity, leading to more confident genetic instruments for gene expression.

Despite the improved performance in predicting gene expression attributable to models with multiple, independent SNPs, we found that, using available GTEx data, TWAS associations based on these models were, in most instances, driven by a single SNP within each trait-associated locus: 90% of associations were no longer significant after stepwise conditional analysis. Although this proportion is likely to fall as eQTL sample sizes increase (increasing the power to detect the additional impact of conditioned variants), these results indicate that, for many genes, the increment in power gained by moving from single to multi-SNP analyses is modest.

The genetic architecture underlying metabolite traits provides a unique opportunity to quantify the performance of gene associations based on multi-SNP models. By leveraging a “truth” set of experimentally validated genes linked to metabolites, we have shown, using GTEx, that TWAS has reasonable sensitivity (67%) at identifying causal genes. However, the PPV is low (8%), as a great majority of associations in the vicinity of a known causal gene involved nearby “bystander” genes. Furthermore, the process of resolving true causal from false-positive associations is complicated by the fact that these types of associations were indistinguishable in their model SNP effect sizes (GWAS and eQTL), colocalization probabilities, and distance to transcription start sites. In the case of the metabolite glutarylcarnitine, for example, the met-QTL rs8012 not only regulates the expression of the causal *GCDH* gene but also five other genes at the same locus, all of which are associated with glutarylcarnitine levels in TWAS. These insights temper the extent to which it can be assumed that genes implicated by significant TWAS associations are causal.

These “bystander” effects reflect their shared regulatory architecture with known causal genes, and our observations around met-QTLs mirror recent findings at the *SORT1* and *NOD2* loci (associated with LDL cholesterol and Crohn’s disease, respectively)^33^. By anchoring our analysis on a wide range of metabolomic phenotypes, we have been able to extend those observations, and to develop more generalizable estimates of the sensitivity and PPV of TWAS. Recent analyses from Stacey and colleagues using an alternative gene prioritisation method (ProGeM) are also instructive^17^. Using ProGEM to address a similar problem (the detection of causal effector genes at met-QTL loci), the performance was appreciably better than that we observed with a sensitivity of 98%, and a specificity ranging from 38.4% to 84.6% (PPV was not measured, and the true negatives needed for estimation of specificity were derived using different criteria for delineating sets of candidate causal genes). However, in contrast to TWAS, ProGeM explicitly integrates SNP-level annotations (i.e. eQTLs) with functional gene and pathway annotations across five databases to prioritize causal genes. That is to say, ProGem directly leverages molecular pathway annotations whereas TWAS is agnostic to this information. Accordingly, ProGeM is intended for a specific trait class - molecular QTLs (e.g. metabolites, lipids, proteins) - and the incorporation of additional information relevant to metabolites is likely to have contributed to the better performance in this specific task. In addition, the sensitivity of ProGeM may be inflated by the fact that shared database features were used to both prioritise genes and benchmark performance. For these reasons, ProGEM might be expected not to achieve comparable performance when used to prioritise effectors at disease GWAS loci, with performance more resembling that of the more agnostic approach we achieve with TWAS.

Our analyses were focused on the use of expression QTLs to map causal genes at metabolomic-QTL signals: the extent to which similar observations apply to other molecular QTLs remains to be determined. Previous studies have shown that the genetic architecture of protein-QTLs (pQTLs) is distinct from that of eQTLs: only half of pQTLs identified in lymphoblastoid cell lines (LCLs) were also eQTLs, and pQTL effect sizes were typically lower than those for eQTLs^34^. However, these apparently distinct architectures are likely in part the consequence of disparities in sample sizes and differences in the technologies used to profile these features. Further work is required to assess if the confounding effect of co-regulation observed in TWAS based on predicted gene expression will be present to the same extent for other molecular features.

TWAS approaches provide an attractive option for prioritizing candidate genes at trait-associated loci. Here, we have demonstrated the potential for these approaches to identify associations that are not causal, through a combination of incomplete colocalization and pleiotropy in gene expression regulation. Ultimately, the process of identifying causal genes at GWAS signals represents an integrative enterprise that is dependent on combining results from multiple complementary approaches, including, in addition to QTL-mapping, epigenome profiling (e.g. chromatin co-accessibility or conformation capture methods), functional screens (e.g. high-throughput gene knock-out CRISPR screens) and the detection of coding variant associations. All of these prioritization approaches – including TWAS – will become more accurate, as the data sets available encompass a wider range of tissues and cell types captured in circumstances (e.g. developmental stages, physiological states, environmental exposures) that better reflect the underlying pathophysiology of the particular traits and diseases under investigation.

## Supplemental Data

Supplemental Figures. Figures S1 and S2

Supplemental Tables. Tables S1-S8

## Acknowledgments

MMcC is a Wellcome Investigator and an NIHR Senior Investigator. Relevant funding support for this work comes from Wellcome (090532, 106130, 098381, 203141, and 212259), NIDDK (U01-DK105535; U01-DK085545) and NIHR (NF-SI-0617-10090). AP was supported by the Rhodes Trust, the Natural Sciences and Engineering Research Council of Canada, and the Canadian Centennial Scholarship Fund. While employed at the University of Oxford, MvdB was supported by a Novo Nordisk postdoctoral fellowship run in partnership with the University of Oxford. This work was also supported by Oxford Biomedical Research Computing (BMRC) facility, a joint development between the Wellcome Centre for Human Genetics and the Big Data Institute supported by Health Data Research UK and the NIHR Oxford Biomedical Research Centre. The views expressed are those of the author and not necessarily those of the NHS, the NIHR or the Department of Health.

## Web Resources

The pre-trained multi-SNP models across 43 GTEx (version 7) tissues are available at http://mccarthy.well.ox.ac.uk/pub/

