## Supplemental Figures for "A multi-tissue transcriptome analysis of human metabolites guides the interpretability of associations based on multi-SNP models for gene expression"

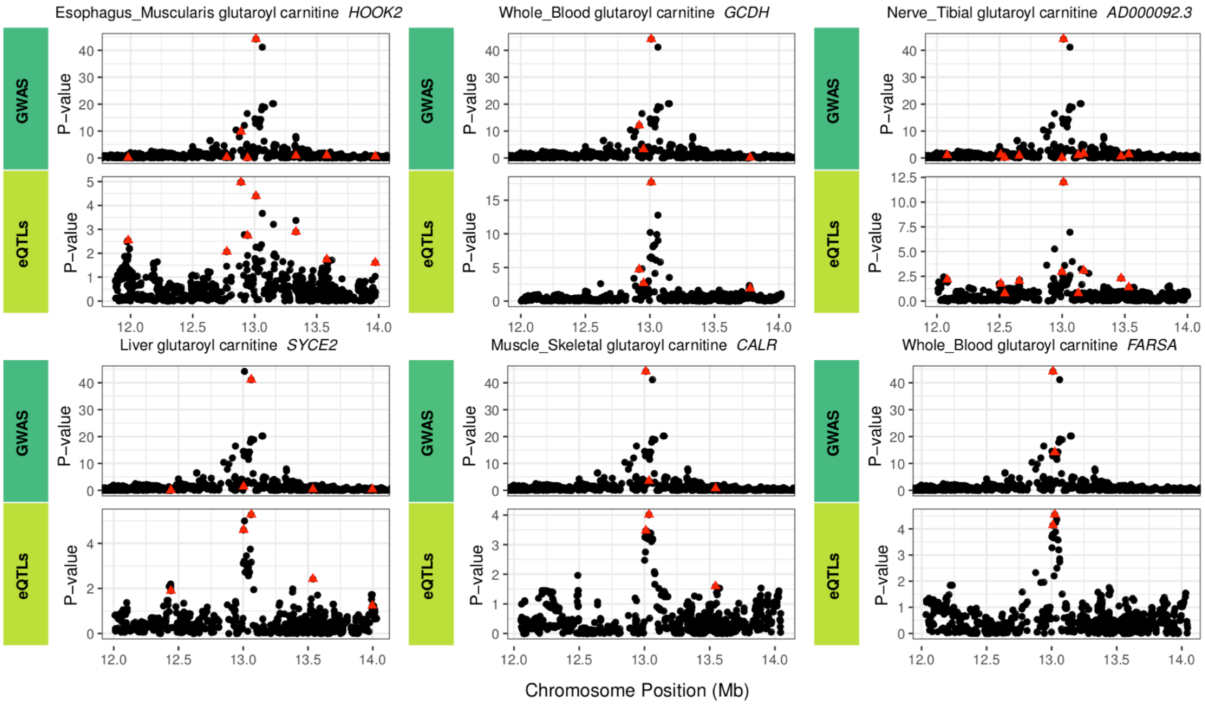


**Figure S1. Gene expression co-regulation at the *GCDH* locus.** Six genes at the *GCDH* locus whose predicted expression significantly mediated glutarylcarnitine levels. The multi-SNP models for all six genes included the *cis*-eQTL rs8012 that regulates the expression for all the genes. *GCDH* is the true causal gene influencing metabolite levels at this locus.

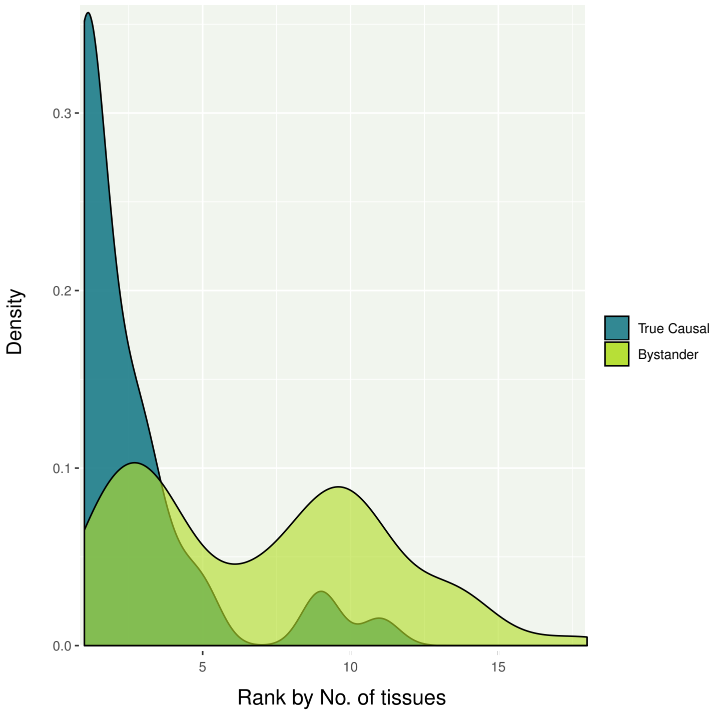


**Figure S2. Causal and bystander gene ranks by number of tissues**. Comparison of ranks by tissue counts for true causal genes and bystander genes at each locus. True causal genes ranked higher (significant in a more tissues) than bystander genes at the same locus.
